# The Influence Of Social Behavior On Competition Between Virulent Pathogen Strains

**DOI:** 10.1101/293936

**Authors:** Joe Pharaon, Chris T. Bauch

**Affiliations:** Dept. of Applied Mathematics, University of Waterloo, 200 University Ave West, Waterloo ON N2L 3G1

## Abstract

Infectious disease interventions like contact precautions and vaccination have proven effective in disease control and elimination. The priority given to interventions can depend strongly on how virulent the pathogen is, and interventions may also depend partly for their success on social processes that respond adaptively to disease dynamics. However, mathematical models of competition between pathogen strains with differing natural history profiles typically assume that human behaviour is fixed. Here, our objective is to model the influence of social behaviour on the competition between pathogen strains with differing virulence. We couple a compartmental Susceptible-Infectious-Recovered model for a resident pathogen strain and a mutant strain with higher virulence, with a differential equation of a population where individuals learn to adopt protective behaviour from others according to the prevalence of infection of the two strains and the perceived severity of the respective strains in the population. We perform invasion analysis, time series analysis and phase plane analysis to show that perceived severities of pathogen strains and the efficacy of infection control against them can greatly impact the invasion of more virulent strain. We demonstrate that adaptive social behaviour enables invasion of the mutant strain under plausible epidemiological scenarios, even when the mutant strain has a lower basic reproductive number than the resident strain. Surprisingly, in some situations, increasing the perceived severity of the resident strain can facilitate invasion of the more virulent mutant strain. Our results demonstrate that for certain applications, it may be necessary to include adaptive social behaviour in models of the emergence of virulent pathogens, so that the models can better assist public health efforts to control infectious diseases.

## 2 Introduction

Modern approaches to developing a theory of the spread of infectious diseases can be traced to 1927 when Kermack and McKendrick developed an integro-differential equation model now widely described as the SIR (Susceptible-Infected-Recovered) model [1]. The model tracks changes in the number of individuals susceptible to an infection *S*(*t*), the number of infected individuals *I*(*t*), and (implicitly) the number of recovered individuals *R*(*t*). Compartmental models such as the SIR model are useful for mechanistic modelling of infection transmission in populations. They have since been further developed to study the evolution and epidemiology of multiple species of pathogens in a population or different strains of the same species [2]. Some models focus on between-host competition while some others on within-host competition [3]. Bull suggested in the 1990s that coupling inter-host and intra-host dynamics in models may be desirable [4]. Models linking between-host transmission dynamics to within-host pathogen growth and immune response are now becoming commonplace [3, 5, 6, 7, 8]. One such approach is to link host viral load (which is a necessary condition of virulence) to the between-host transmission rate.

Compartmental models have also been used to study the phenomenon of pathogen virulence–the rate at which a pathogen induces host mortality and/or reduces host fecundity [9, 10, 11, 12]. It was initially believed that hosts and parasites co-evolved to a state of commensalism (whereby parasites benefit from their host without harming them) [13, 14] but this hypothesis was later challenged [15]. In mathematical models, virulence is often treated as a fixed model parameter expressing the excess mortality rate caused by the pathogen. For instance, virulence has been assumed to depend on the intrinsic reproductive rate of the parasite [16]. Other research expresses the transmission rate *β* and the recovery rate *μ* in terms of a parameter *ν* that represents virulence [17]. When the impact of human behaviour is discussed in such models, it is discussed in terms of hypothesized effects of human behaviour on the value of the fixed parameter representing virulence. A Human Immunodeficiency Virus (HIV) virulence model by Massad *et al* [18] shows that reducing the number of sexual partners could possibly drive HIV to be a more benign pathogen. However, the model assumes that the number of sexual partners can simply be moved up or down as a model parameter, whereas in reality the number of sexual partners in a population is the outcome of a dynamic socio-epidemiological process that merits its own mechanistic modelling, and itself responds to pathogen virulence. In general, these models do not treat human behaviour as a dynamic variable that can evolve in response to transmission dynamics and influence the evolution of virulence. (A few exceptions exist, including work that allows virulence to be a function of the number of infected hosts, thus capturing a situation where the magnitude of the epidemics affects the ability of health care services to host patients [19].) However, as human responses to both endemic and emerging infectious diseases show, human behaviour can have a significant influence on how infections get transmitted. For instance, an early and well-documented example shows how the residents of the village of Eyam, England quarantined themselves to prevent the spread of plague to neighbouring villages [20]. Individuals moved to less populated areas during the Spanish Influenza pandemic in the early 20th century [21]. More recently, masks became widely used during the outbreak of the Severe Acute Respiratory Syndrome (SARS) at the beginning of the 21st century [22], and it has been shown pathogen virulence in Marek’s disease can evolve in response to how vaccines are used [23].

Theoretical models of the interactions between human behaviour and the spread of infectious diseases are increasingly studied [24, 25, 26, 27, 28, 29, 30]. For instance, Bagnoli *et al* [31] found that under certain conditions, a disease can be driven extinct by reducing the fraction of the infected neighbours of an individual. Zanette *et al* [32] showed that if susceptible individuals decide to break their links with infected agents and reconnect at a later time, then the infection is suppressed. Gross [33] also shows that rewiring of edges in a network (and thus social interaction) can greatly influence the spread of infectious diseases. Of the compartmental models, we focus on those that have used concepts from evolutionary game theory such as imitation dynamics [34] to describe the evolution of behaviour and its interplay with the epidemics. An example of imitation dynamics concerns, as described in detail in [35], the effect of vaccination on the spread of infectious diseases. Each individual in the population picks one strategy and adopts it: “to vaccinate” or “not to vaccinate”. The proportion of vaccinators is modelled using an ordinary differential equation and is coupled with a standard SIR model. An important aspect of behavioural models is to couple them with epidemiological processes such as transmission. This coupling creates a feedback loop between behaviour and spread of the disease.

Given that adaptive social behaviour is important in many aspects of infection transmission, we hypothesize that adaptive social behaviour can also influence selection between pathogen strains with differing virulence in ways that cannot be captured by assuming it to be represented by a fixed parameter. Our objective in this paper was to explore how behaviour and virulence influence one another, in a coupled behaviour-disease differential equation model. The model allows individuals who perceive an increase in the prevalence of infection to increase their usage of practices that reduce transmission rates (such as social distancing and hand-washing) and thereby boost population-level immunity. This approach can help us understand the effects specific social dimensions, such as level of concern for a strain or the rate of social learning, on the coupled dynamics of pathogen strain emergence and human behaviour in a situation where virulence imposes evolutionary trade-offs and is strain-specific. Instead of considering long-term evolutionary processes with repeated rounds of mutation and selection, we focus on the case of invasion of a single mutant strain with a large phenotypical difference compared to the resident strain. In the next section, we describe a model without adaptive social behaviour as well as a model that includes it, and in the following Results section we will compare their dynamics.

## 3 Model

We compare dynamics of a two-strain compartmental epidemic model in the presence and absence of adaptive social behaviour. Individuals are born susceptible (*S*). They may be infected either by a resident strain (*I*_1_) or a mutant strain (*I*_2_). For simplicity, we assume that co-infection and super-infection are not possible. Infected individuals can either recover (*R*) or die from infection. We furthermore assume that recovery from either strain offers permanent immunity to both strains. The system of differential equations representing the *SI*_1_*I*_2_*R* model in the absence of adaptive social behaviour (we will refer to this as the “uncoupled model” throughout) is given by

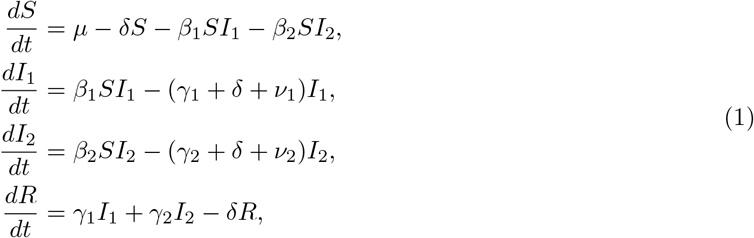

where *β*_1_ (*β*_2_) represents the transmission rate of the resident (mutant) strain; *γ*_1_ (*γ*_2_) represents the recovery rate from the resident (mutant) strain; *ν*_1_ (*ν*_2_) represents the death rate from the resident (mutant) strain due to infection (virulence); *μ* is a birth rate and *δ* is the background death rate. All variables represent the number of individuals with the given infection status (for instance, *S* is the number of susceptible individuals). Since *R* does not appear in the other equations, we can omit *R* from the analysis.

The system of differential equations in the presence of adaptive human behaviour couples the *SI*_1_*I*_2_*R* epidemic spread with a differential equation for human behaviour (“coupled model”). Each individual in the population can choose to accept or reject behaviours that reduce infection risk (e.g. washing hands, wearing a mask, social distancing), and individuals imitate successful strategies observed in others. Let x represent the proportion of individuals accepting preventive behaviour (we will call these “protectors”). Individuals sample others in the population at rate κ, representing social learning. The choice is based on the perceived severity *ω*_1_ (resp. *ω*_2_) from the resident (resp. mutant) strain, where *ω*_1_ (resp. *ω*_2_) can be quantified as the probability that an infection by the resident (resp. mutant) strain results in a severe case of disease. The more severe cases the population observes, the more attractive preventive behaviour becomes: we assume that individuals respond to the total number of severe cases *ω*_1_*I*_1_*I*_2_ + *ω*_2_*I*_2_ they observe at a given time. Preventive behaviour is not always completely effective. We introduce efficacy of infection control *ϵ*_1_ (*ϵ*_2_) against the resident (mutant) strain. The efficacy of infection control influences the transmission process. The more effective infection control is against a strain, the less likely it will be transmitted.

More formally, the preceding imitation dynamic (or equivalently, replicator dynamic) assumes that each individual samples others as a fixed rate, and if another person is found to be playing a different strategy but is receiving a higher payoff, the individual switches to their strategy with a probability proportional to the expected gain in payoff [36]. These assumptions give rise to a differential equation of the form *dx/dt* = *kx*(1 − *x*)Δ*U* where *k* is the sampling rate and Δ*U* is the payoff difference between the two strategies. This equation is derived elsewhere and is used in other socio-ecological and socio-epidemiological models [35, 37, 38, 39]. The augmented system of differential equations representing the coupled social-epidemiological *SI*_1_*I*_2_*RX* model with adaptive human behaviour is therefore given by:

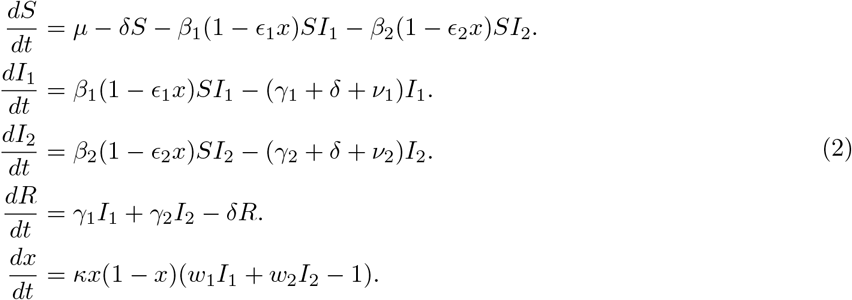

We apply the restrictions *ϵ_i_* ∈ [0,1] and *ω_i_* ≥ 0.

Baseline parameter values are summarized in Table 1. We chose parameter values to represent an emerging infectious disease with a relatively low basic reproduction number and an acute-self limited infection natural history, as might occur for viral infections such as ebola or influenza. Recruitment is assumed to occur due to births and immigration at a constant rate *μ*, while the per capita death rate due to all causes other than the infection is *δ*. The values of *μ* and *δ* are obtained as the reciprocal of an average human lifespan of 50 years. Note that *γ_i_* + *ν_i_* is the reciprocal of the average time spent in the infected class before the individual recovers or dies from infection. Since we are assuming that strain 2 is more virulent, *ν*_2_ − *ν*_1_ = 0.05/day can be considered as the excess death rate due to infection from the more virulent strain 2. We assume *β*_1_ = *β*_2_ and therefore 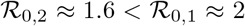. Hence, all else being equal, the more virulent strain has a lower reproductive number and is therefore at a disadvantage to invade. We note that *R* does not appear in the other equations and hence could be omitted.

**Table 1:** Baseline parameter values. Strain 1 is taken to be an avirulent resident strain, and strain 2 is taken to be a more virulent mutant strain.

| Parameter | Definition | Value |
| --- | --- | --- |
| $\delta$ | death rate | 1/18250 per day, [34]. |
| $\mu$ | birth rate | 1/18250 per day, [34]. |
| $\gamma_1$ | recovery rate for strain 1 | 0.2 per day (assumed). |
| $\nu_1$ | disease death rate for strain 1 | 0.0 per day (assumed). |
| $\gamma_2$ | recovery rate for strain 2 | 0.2 per day (assumed). |
| $\nu_2$ | disease death rate for strain 2 | 0.05 per day (assumed). |
| $\beta_1$ | transmission rate for strain 1 | 0.4 per day (assumed). |
| $\beta_2$ | transmission rate for strain 2 | 0.4 per day (assumed.) |
| $\kappa$ | sampling rate | 1/365 per day, [37]. |
| $w_1$ | perceived severity from strain 1 | 10000 (assumed) |
| $w_2$ | perceived severity from strain 2 | 100000 (assumed) |
| $\epsilon_1$ | efficacy of infection control against strain 1 | 0.7 (assumed) |
| $\epsilon_1$ | efficacy of infection control against strain 2 | 0.4 (assumed) |

We identify all equilibria of the uncoupled and coupled systems and determine their local stability properties. We study conditions under which the mutant strain successfully invades. Due to the analytical complexity of the coupled model, we rely primarily on numerical simulations. We used MATLAB to run our simulations and generate parameter planes (ODE45, ODE23tb, and ODE15s). We also wrote MATLAB code to analyze the stable regions of all equilibria versus a combination of parameters of interest.

## 4 Results

### 4.1 Invasion analysis: *SI*_1_*I*_2_*R* model

The *SI*_1_*I*_2_*R* model has 3 equilibria [40]. One equilibrium is disease free and is stable when

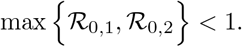

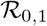 (resp. 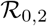) is the basic reproductive number of the resident (mutant) strain, where *R*_0, *i*_ is given by

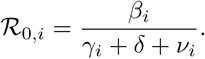

The other two equilibria are endemic. Assuming that basic reproductive numbers are not equal, strains can not co-exist and the strain with the higher basic reproductive number always invades.

### 4.2 Invasion analysis: *SI*_1_*I*_2_*RX* model

In contrast, the addition of a dynamic social variable x(*t*) generates 9 equilibria for the *SI*_1_*I*_2_*RX* model. Two equilibria are disease free and the other equilibria are endemic. One of the 7 endemic equilibria represents a state of coexistence of both strains (which can occur even if basic reproductive numbers are not the same). The analytical expression and stability criteria for the equilibrium with co-existing strains are difficult to compute and therefore, we analyze it numerically.

If both basic reproductive numbers are less than 1, then the system is disease-free and social behaviour is not relevant. Assume, on the other hand, that 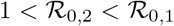 (as in our baseline parameter values, Table 1, and where the expressions for 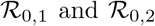 are the same as in the *SI*_1_*I*_2_*R* model and assume *x* = 0. This corresponds to a scenario where the resident strain is more transmissible than the mutant strain. As already noted the mutant strain can not invade in the absence of adaptive social behaviour (*SI*_1_*I*_2_*R* model). However, in the presence of adaptive social behaviour, we can derive necessary and sufficient conditions for the mutant strain to invade when 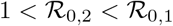:

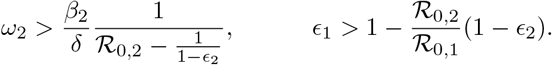

These results show that a high level of perceived severity from the mutant strain is a necessary condition for invasion. However, it has to be coupled with a sufficiently high efficacy of infection control against the resident strain (with *ϵ*_1_ > *ϵ*_2_). A high efficacy of infection control against the resident strain will effectively reduce its transmission and therefore creating a larger pool of susceptible individuals for the mutant strain. The two conditions must be met simultaneously to provide a necessary and sufficient condition for invasion. The condition that *ϵ*_1_ > *ϵ*_2_ could easily be met in a real population if the two strains differ in their model of transmission, and the population has more experience with controlling the resident strain than with the new mutant strain. Moreover, a high value of *ω*_2_ could easily be met in a real population due to spreading panic about a new and more virulent strain that public health does not yet know how to best control.

We also consider necessary and sufficient conditions for failure of invasion of the mutant strain:

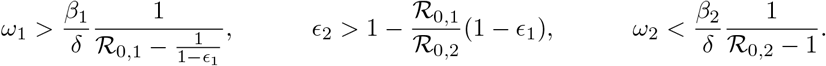

Invasion fails when perceived severity of the mutant strain is low enough but also that of the resident strain high enough. Note the difference between invasion and failure to invade. Here, we require conditions on both perceived severities. As predicted, if the efficacy of infection control against the mutant strain is high enough (relative to that of the resident strain) then invasion fails. Again, all three conditions must be met jointly. Together, they create a necessary and sufficient condition for the failure of invasion.

Finally assume that 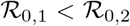 (this scenario is not discussed at length in this paper). In the absence of social behaviour, the mutant strain is bound to invade. However, we derive necessary and sufficient conditions for the failure of invasion when social behaviour is added to the system:

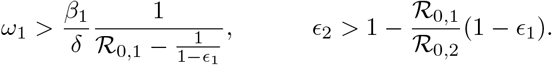

Note the difference between this case and the case when 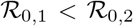: there is no conditioning on *ω*_2_. If the mutant strain has a higher fitness, it does not matter how severely it is perceived (since individuals respond to the weighted sum of mutant and resident prevalence and the mutant is initially rare, hence the early response is dominated by the resident). It will fail to invade provided that the perceived severity of the resident strain is high enough and that efficacy of infection control against the mutant strain is high enough relative to the resident strain (with *ϵ*_2_ > *ϵ*_1_). Once again we require both inequalities to be satisfied and together they provide necessary and sufficient condition for the mutant strain to fail invasion.

We finally turn our attention to the invasion of the mutant strain when it is more transmissible. The invasion is conditional: necessary and sufficient conditions for the invasion of the mutant strain are:

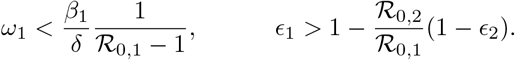

In this scenario, a low perceived severity of the resident strain will allow invasion of the mutant strain provided that the efficacy of infection control against the resident strain is high enough.

The addition of adaptive social behaviour to the epidemic model introduced four new parameters, and it is clear that the model permits conditions for the mutant strain to invade on account of behaviour, even when the mutant strain has a lower basic reproductive ratio, as long as certain conditions for efficacy of infection control are satisfied and level of concern about the severity of the mutant strain are satisfied. To gain further insight into how adaptive social behaviour influences the invasion of the more virulent strain, we turn to numerical simulation and generation of time series and parameter planes.

### 4.3 Time series analysis

We use time series of model simulations to illustrate some of the model’s dynamical regimes. We consider the case where *ν*_1_ =0 and *ν*_2_ = 0.25 and therefore 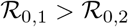 while assuming (for simplicity) that *β*_1_ = *β*_2_ and *μ*_1_ = *μ*_2_. Hence, the mutant strain is more virulent and kills its hosts more quickly, giving it a significantly lower basic reproduction number. We use a simulation time horizon on the order of hundreds of years-although both pathogen and social parameters could vary over this period, a long time horizon ensures that the asymptotic model states are correctly characterized, and thus enables us to meet our objective of gaining insight into the types of dynamics exhibited by the model.

We first consider a scenario where the mutant strain, on account of its greater virulence, is perceived to be ten times more severe than the resident strain (*ω*_2_ = 10^5^ = 10*ω*_1_). Moreover, infection control against the resident strain is much more effective, on account of less being known about the modes of transmission of the mutant strain (baseline values: *ϵ*_1_ = 0.7 > *ϵ*_2_ = 0.4). In this scenario, the mutant strain invades (Figure 1a). This agrees with the conditions determined in our invasion analysis. We observe that the mutant strain quickly displaces the resident strain and converges to an endemic state where the proportion of protectors *x* remains relatively high (Figure 1a). On shorter timescales, we see a transient phase at the start of the simulation with a sharp epidemic of the resident strain, followed by periodic epidemics with much lower incidence of the mutant strain (Figure 1b,c). The numerical simulations agree with the values computed from analytical expressions at equilibrium (for sufficiently large values of *t*).

**Figure 1:**
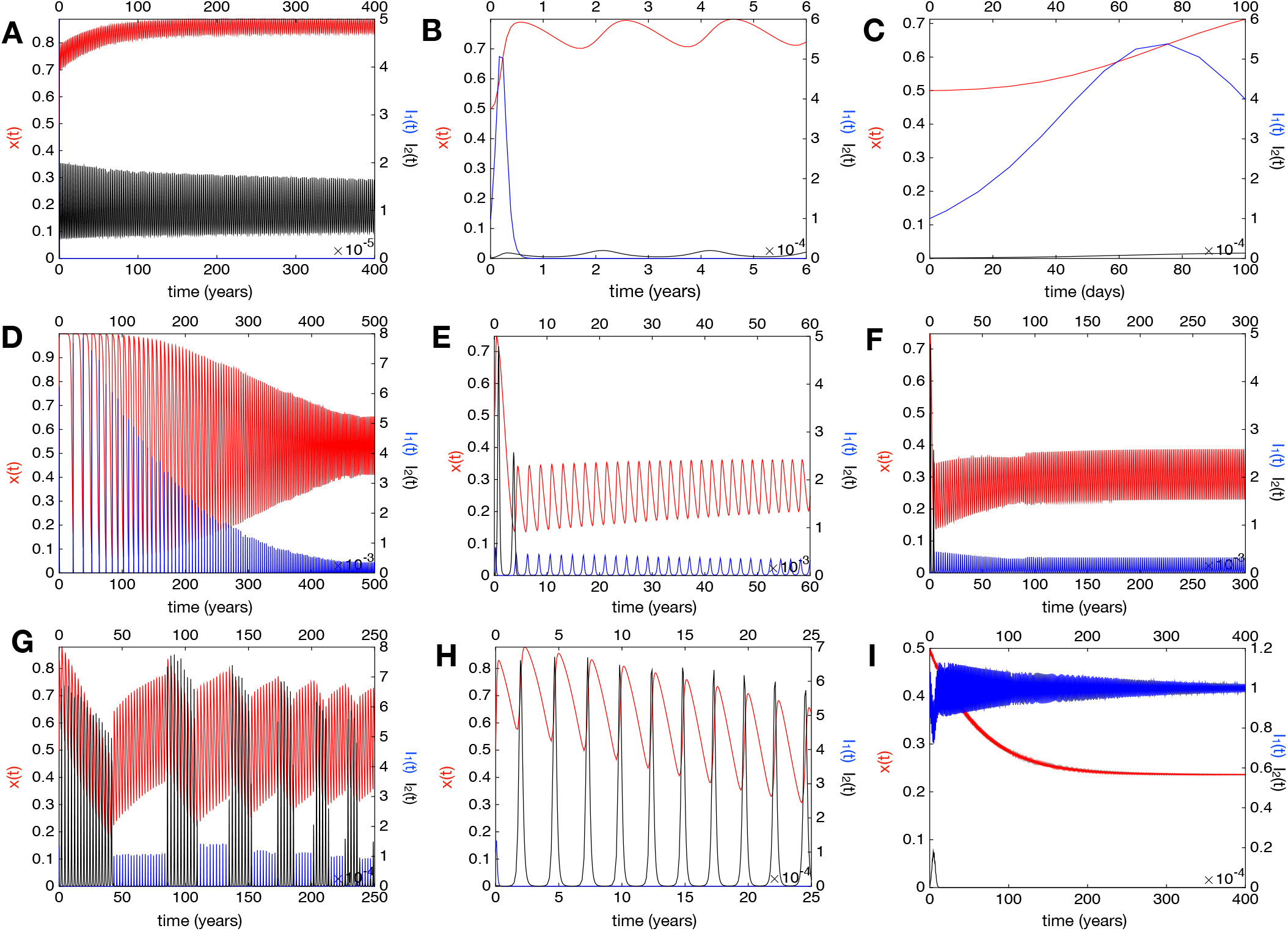
Numerical simulations for the *SI*_1_*I*_2_*RX* model at various values for the social and infection control parameters. (**a,b,c**) show baseline values where the mutant strain is perceived to be 10 times more severe (*ω*_2_ = 10*ω*_1_ = 10^5^) and where efficacy of infection control against the resident strain is greater *ϵ*_1_ = 0.7 > *ϵ*2 = 0.4. The dynamics are shown at different timescales in (**a**), (**b**) and (**c**). (**d**) *ϵ*1 = 0.4. (**e,f**) *ω*_2_ = 10^2^, *ϵ*_2_ = 0.3. (**g,h**) *ω*_1_ = 10*ω*_2_ = 10^5^. (**i**) *ω*_1_ = *ω*_2_ = 10^4^. *ϵ*_1_ = 0.9, *ϵ*_2_ = 0.6. All other parameters are held at their baseline values. Red line represents prevalence of protectors *x*. Blue line represents prevalence of the resident strain *I*_1_. Black line represents prevalence of the more virulent mutant strain *I*_2_.

Decreasing the efficacy of infection control against the resident strain and equating it to that of the more virulent strain (*ϵ*_1_ = *ϵ*_2_ = 0.4, with all other parameter values at baseline values) prevents the invasion of the mutant strain (Fig. 1d). This occurs because more susceptible individuals will be infected by the resident strain, thereby significantly decreasing the pool of susceptible individuals available for infection by the mutant strain.

A surprising scenario under which the invasion of the mutant strain fails is when both perceived severity of the mutant strain and the efficacy of infection control against are low (Figure 1e,f, *ω*_2_ = 10^2^ and e_2_ = 0.3 with other parameter values at baseline). It is worth noting in this case that we initially have a few outbreaks of the mutant strain with high incidence. Fig 1f represents the same dynamics as Fig 1e but on a longer time scale. The oscillations in the prevalence of infection and the prevalence of protectors is typical of coupled behaviour-disease models with adaptive social behaviour [35].

It is difficult for both strains to co-exist without imposing *ω*_1_ = 10^5^ > *ω*_2_ = 10^4^. If the resident strain is perceived to be ten times more severe, then co-existence is achieved via a transient but very long-term pattern of switching between oscillatory regimes before the system finally converges to an equilibrium of co-existence (Fig 1g). The system switches between a longer-lived regime with relatively small epidemics of the resident strain, and a shorter-lived regime with very large epidemics of the mutant strain. Changes in the proportion adopting contact precautions, *x*, facilitates the switching. As *x* rises, it allows a series of periodic outbreaks of the mutant strain which in turn decreases the proportion of people adopting prevention and starts a series of outbreaks of the resident strain. This loop continues with diminishing switching-period as well as amplitude. If we bring back efficacies of infection control to baseline values, this phenomenon persists but with wilder oscillations in *x*. This happens because lower values of *ϵ* increase the effective transmission rate which in turns leads to rapid changes in *x*. Figure 1h shows the same dynamics as in Figure 1g but on a shorter timescale.

We also allowed the perceived severities to be equally high (*ω*_1_ = *ω*_2_ = 10^4^) and we have increased the efficacies from their baseline values (*ϵ*_1_ = 0.9 and *ϵ*_2_ = 0.6) (Figure 1i). We observe that the mutant strain fails to invade and the prevalence of the resident strain remains relatively close to the initial condition.

In order to refine our understanding of the influence of social parameters on the invasion of the mutant strain, we proceed in the next subsection with phase plane analysis that studies the interplay between the parameters determining regions of invasion.

### 4.4 Phase plane analysis

Surprisingly, there are parameter regimes where increasing the perceived severity of the resident strain (*ω*_1_) allows the mutant strain to invade (Figure 2a-c). This occurs across a nontrivial portion of parameter space despite the fact that 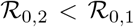. This regime shift occurs because a sufficiently high perceived severity of the resident strain creates a large pool of susceptible individuals, and coupled with a higher efficacy of infection control against the resident strain, this means that the invading mutant strain can take advantage of the increased pool of susceptible individuals to invade. This effect occurs only when the efficacy of infection control against the resident strain is relatively high (e.g. *ϵ*_1_ = 0.9 and *ϵ*_2_ = 0.6). However, this phenomenon does not hold when *ϵ*_1_ and *ϵ*_2_ are low, in which event the model behaves similar to the *SI*_1_*I*_2_*R* model where the strain with the higher basic reproductive number invades, as expected. Similarly, increasing *ω*_2_ can push the system from a regime of co-existence of the two strains to a region where only strain 2 persists.

**Figure 2:**
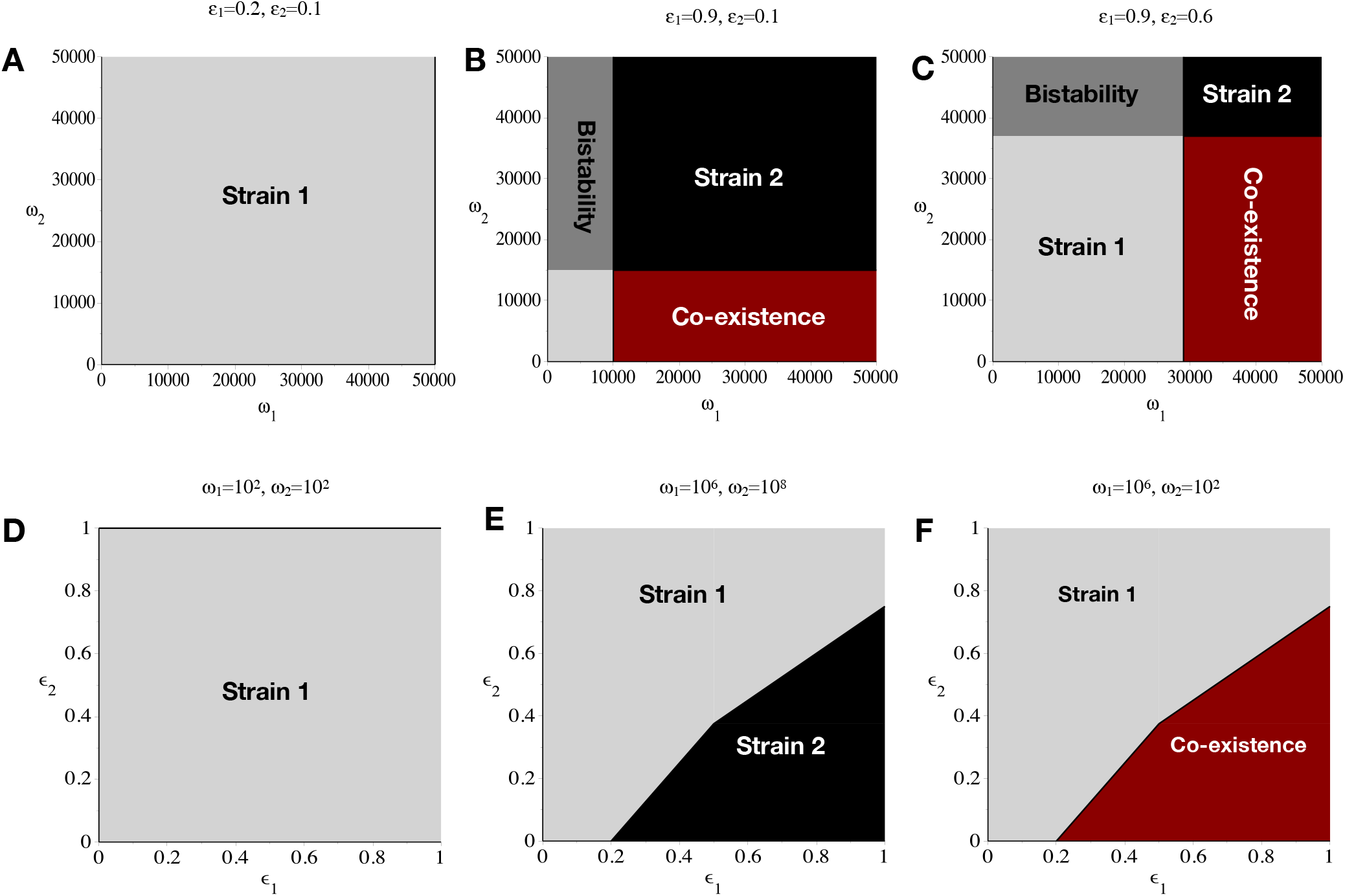
Parameter plane analysis of the *SI*_1_*I*_2_*RX* model. These dynamics are more complex than those exhibited by the *SI*_2_*I*_2_*R* model, which only predicts persistence of strain 1 for equivalent parameter values. The epidemiological parameters are at baseline values (Table 1). The social parameters are varied. (**a**) and (**d**) show no invasion of the mutant strain when *ϵ*_1_ = 0.2 and *ϵ*_2_ =0.1 in the *ω*_1_ − *ω*_2_ parameter plane (**a**) and when *ω*_1_ = *ω*_2_ = 10^2^ in the *ϵ*_1_ — *ϵ*_2_ parameter plane (**d**). (**b**) and (**c**) represent similar qualitative results when for large *ϵ*_1_ = 0.9 we get invasion of the mutant strain in the black region and co-existence with the resident strain in the red region. The invasion region is bigger when *ϵ*_2_ is lower (e_2_ = 0.1 in (**b**) and *ϵ*_2_ = 0.6 in (**c**)). Finally, in (**e**) and (**f**) we observe qualitatively different results when we vary **ω*_2_* in the *ϵ*_1_ − *ϵ*_2_ parameter plane. In (**e**), 10^8^ = *ω*_2_ > *ω*_1_, we have invasion of the mutant strain. In (**f**), *ω*_1_ = 10^6^ > *ω*_2_, we have co-existence of the strains. The light gray region in the lower-left hand corner of subpanel (**b**) corresponds to both strains being extinct.

In *ϵ*_1_ = *ϵ*_2_ parameter planes we again find parameter regimes where the more virulent strain can invade due to adaptive social behaviour, despite the fact that 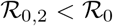, if there is an imbalance in the perceived severity of the two strains. When perceived severities are sufficiently low, the mutant strain can never invade (Figure 2d). But when *ω*_2_ >> *ω*_1_, the mutant strain can invade and remove the resident strain if *ϵ*_1_ is sufficiently large and *ϵ*_2_ is sufficiently small (Figure 2e). When *ω*_1_ >> *ω*_2_, the mutant strain and the resident strain coexist, when *ϵ*_1_ is sufficiently large and *ϵ*_2_ is sufficiently small (Figure 6f). Increasing the efficacy of infection control against the resident strain (*ϵ*_1_) or decreasing efficacy of control against the mutant strain (*ϵ*_2_) can allow the mutant strain to invade (Figure 2e-f). We note again that, surprisingly, invasion can result in the elimination of the resident strain if the perceived severity of the mutant strain is significantly higher than that of the resident strain (*ω*_2_ >> *ω*_1_), but when the opposite applies, coexistence results.

## 5 Discussion

We have showed how adaptive social behaviour greatly impacts the evolution of virulence in a coupled behaviour-disease model. If we neglect social behaviour, the basic reproductive numbers of the two strains are sufficient to predict which of the strains will invade a population. However, adding adaptive social behaviour with asymmetric stimulation and effects on either strain to an epidemiological system completely shifts how we view whether a more virulent strain will be selected for. As we have seen, social behaviour can either act in favour or against the invasion of a more virulent strain, and we can describe these effects with reference to specific social parameters (*ω*_12_) quantifying how concerned individuals are about the two strains, and control parameters (*ϵ*_1,2_) quantifying how well infection control measures like hand-washing work. Most interestingly, adaptive social behaviour enables invasion of the mutant strain under plausible epidemiological and social conditions even when it has a lower basic reproductive number.

Future work can generate further insights into how behaviour and virulence interact for specific infectious diseases, by building on existing research on the coupled dynamics of behaviour and infection transmission. For instance, an increase in the average number of sexual partners of an individual has been predicted by mathematical models to cause increased HIV virulence [18, 41]. These models use a fixed parameter to quantify the number of sexual partners, but the number of sexual partners could be made to evolve dynamically based on the number of infected individuals in a particular population, similar to seminal work using compartmental models to model core group dynamics [42]. An increase in the number of sexual partners will decrease the efficacy of infection control against the more virulent strain and effectively increase its transmission and hence leads to higher virulence. Other future research could explore how adaptive social behaviour interacts with evolutionary trade-offs to determine virulence evolution. One of the most common hypotheses is that a trade-off exists for between-host transmission and virulence. To increase its probability of transmission, the parasite must replicate within the host. This replication, on the other hand, must be controlled because otherwise it might lead to the host’s death and therefore prevent transmission. However, other trade-offs have been suggested, such as between transmission rate and host recovery rate [43]. Moreover, complicated host life cycles imply that many other types of trade-offs are also possible [44], and the presence of multiple trade-offs may complicate the relationship between transmission rate and virulence [45]. Social behaviour could interact with evolutionary trade-offs to alter the virulence evolution of an emerging pathogen, and this process could be modelled by building on existing virulence evolution models.

While the model discussed in this paper serves as a general framework for studying the influence of social behaviour on strain competition and emergence, further research needs to be carried out to understand the interplay between the epidemiological and social parameters. For instance, we did not model virulence evolution explicitly but rather by assuming two strains have already emerged due to mutation and addressing conditions under which the more virulent mutant strain is more fit. Future research could instead model virulence by defining transmission and recovery rates in terms of a virulence parameter, or by using an adaptive dynamics approach. Future research could also explore different possible relationships between the virulence parameters *ν*_1,2_ and the perceived severity parameters *ω*_1,2_, or the interaction between social learning timescales and pathogen evolutionary timescales. We did not study the influence of the social learning parameter *κ* in this paper, but previous research on other socio-ecological and socio-epidemiological systems suggests that the social learning rate can destabilize interior equilibria [39, 38]. A model that accounts for multiple rounds of mutation would enable studying how pathogen evolutionary timescales interact with social learning dynamics. Finally, we assumed no specific relationship between the perceived severity *ω*_1,2_ and the virulence *ν*_1,2_ although a non-trivial relationship certainly exists, and future research could explore possible assumptions for their formal relationship.

In conclusion, our model shows how social behaviour can influence the virulence of emerging strains under plausible parameter regimes when using standard models for social and infection dynamics. When analyzing emerging and re-emerging pathogens and continually evolving infectious diseases such as influenza, it is worthwhile further considering aspects of social behaviour in efforts to mitigate serious threats.

